# Pre-stress of actin cortices is important for the viscoelastic response of living cells

**DOI:** 10.1101/783613

**Authors:** Andrea Cordes, Hannes Witt, Aina Gallemí-Pérez, Bastian Brückner, Florian Grimm, Marian Vache, Tabea Oswald, Daniel Flormann, Franziska Lautenschläger, Marco Tarantola, Andreas Janshoff

## Abstract

Shape, dynamics, and viscoelastic properties of eukaryotic cells are primarily governed by a thin, reversibly cross-linked actomyosin cortex located directly beneath the plasma membrane. We obtain time-dependent rheological responses of fibroblasts and MDCK II cells from deformation-relaxation curves using an atomic force microscope to access the dependence of cortex fluidity on pre-stress. We introduce a viscoelastic model that treats the cell as a composite shell and assumes that relaxation of the cortex follows a power law giving access to cortical pre-stress, area compressibility modulus, and the power law (fluidity) exponent. Cortex fluidity is modulated by interfering with myosin activity. We find that the power law exponent of the cell cortex decreases with increasing intrinsic pre-stress and area compressibility modulus, in accordance with previous finding for isolated actin networks subject to external stress. Extrapolation to zero tension returns the theoretically predicted power law exponent for transiently cross-linked polymer networks. In contrast to the widely used Hertzian mechanics, our model provides viscoelastic parameters independent of indenter geometry and compression velocity.

---

Many cellular processes such as adhesion, motility, growth, and development are tightly associated with the mechanical properties of cells and their environment [1–3]. Vitality and fate of cells are often directly inferred from their elastic properties [4–6]. In search for effective and standardized mechanical phenotyping of living cells, several tools have been developed that permit precise and fast measurements [7]. The response of cells to external deformation is primarily attributed to the viscoelasticity of the cellular cortex [8, 9]. The cortex forms a composite shell consisting of a compliant but contractile actin mesh with a large number of actin-binding proteins coupled to the plasma membrane [10, 11]. The thin actin cortex can be contracted by the action of motor proteins such as myosin II, resulting in a measurable pre-stress that provides resistance against deformation at low strain [8, 12]. It was found that rheological parameters of compliant cells such as the complex shear modulus generally obey a power law dependency *G*\* ∝ *ω^β^* over multiple decades in frequency *ω* [13, 14]. The dimensionless power law coefficient *β* characterizes the degree of fluidity and energy dissipation upon deformation, where *β* = 0 represents an ideal elastic solid and *β* = 1 a Newtonian liquid. Values obtained for the power law exponent of living cells usually range between 0.2 – 0.4 for adherent cells suggesting glassy dynamics [4, 13]. *In vitro* experiments and theory suggest that transiently cross-linked actin networks generate a broad spectrum of relaxation times typical for a power law behavior with *β* = 0.5 below the characteristic frequency (2*π/τ*_off_, *τ*_off_ being the unbinding time of the cross-linker) [15]. It is still unclear why rheological properties found for living cells and those of artificial actin cortices are different. Recently, Mulla *et al.* could show that transient cross-linking of actin filaments combined with external stress lead to lowering of the power law exponent [16]. Our goal is to examine how internal stress changes the viscoelastic properties of living cells. Therefore, we require a viscoelastic model that permits to relate pre-stress of cells to fluidity obtained from deformation-relaxation experiments. Our viscoelastic model of the cortex is based on power law rheology and suitable to describe the time-dependent deformation and relaxation of adherent and suspended cells. Drugs like blebbistatin [17] and calyculin A [18] were administrated to arrest and boost myosin activity, respectively, allowing us to alter the intrinsic pre-stress in a predictable fashion. We found that cortex fluidity is decreased with increasing pre-stress, while extrapolation to zero tension recovers the predicted power law exponent of 0.5 found for transiently cross-linked actin networks [19].

From scrutinizing the cortex thickness and its mesh size (Fig. 1A-C and SI) with scanning electron microscopy and fluorescence microscopy, it is safe to treat the cortex as a two-dimensional continuous material neglecting bending stiffness and area shear modulus [20]. The cortex resists deformation only by its area compressibility modulus and pre-stress. We refer to this model as the *viscoelastic Evans* model throughout the text due to his seminal and initial work on cortex mechanics [20]. Viscoelasticity of the 2D area compressibility modulus is assumed to follow a power law. Minimizing free energy assuming constant volume leads to minimal surfaces of constant curvature. The force *f* balance at the equatorial radius for cells between two parallel plates reads:

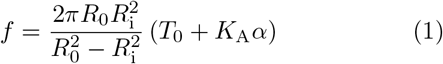

with *R*_0_ the equatorial radius, *R*_i_ the contact radius, 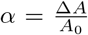 the areal strain, *T*_0_ the pre-stress, and *K*_A_ the area compressibility modulus of the cortex. It is straight-forward to cast the model into non-dimensional form that permits to write eq. (1) as 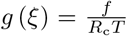, with *ξ* = *z*_p_*/R*_c_ (Fig. 1A, SI). *z*_p_ is the distance between the plates, *R*_c_ the initial radius of the cell in suspension and *T* denotes the overall homogeneous tension. Hence, *g*(*ξ*) and *α*(*ξ*) are generic functions that only need to be computed once. Both functions can easily be approximated by polynomials 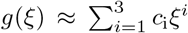 and 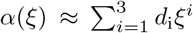 permitting one to obtain an analytical solution of the corresponding elastic-viscoelastic problem (see SI). The general hereditary integral for the restoring force during parallel-plate compression (0 < *t* < *t*_m_, eq. (2)) and relaxation (*t* > *t*_m_) reads [21]:

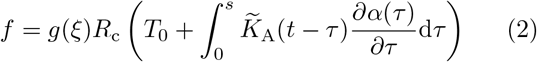

with *s* = *t* for compression and *s* = *t*_m_ for relaxation 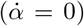. The integrals can be solved by using 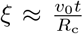 for compression with the constant velocity *v*_0_ and assuming a power law behavior of 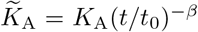 with the time-scaling parameter *t*_0_ (see SI). The general scheme described here can also be used to describe the deformation of adherent and confluent cells with various indenter geometries (see SI).

**FIG. 1.**
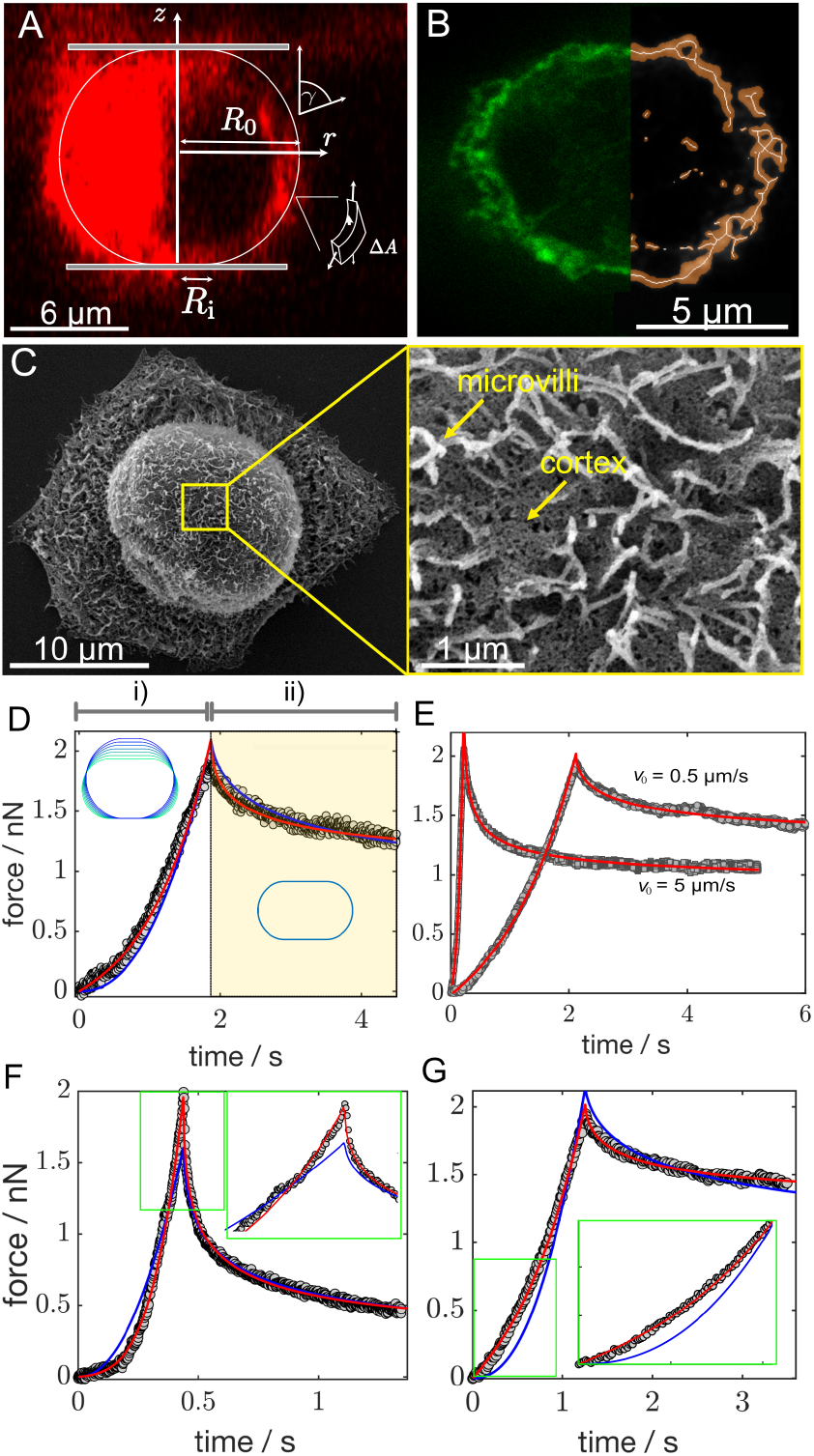
A) CLSM image (*xz* plane) of a MDCK II cell (internal and plasma membrane stained with CellMask) clamped between cantilever and substrate. White drawing shows the parametrization of the cell’s contour. B) Left side shows the STED image (*xy* plane) of the cellular cortex (green: actin) of a MDCK II cell. The right side shows the reconstruction used to determine cortex thickness (see SI). C) SEM image of a MDCK II cell revealing its cortex structure (right). D) Typical compression (i) at *v*_0_ = 0.5 μm followed by force relaxation (ii) of a MDCK II cell and fits according to the Hertz model (blue line, *E*_0_ = 450 Pa, *β* = 0.22) and the Evans model (red line, *T*_0_ = 0.75 mN/m, *K*_A_ = 0.44 N/m, *β* = 0.49). The inset shows the time-evolution of the contour. E) Varying the compression velocity does not significantly change the fitting results: *T*_0_ = 0.83 mN/m, *K*_A_ = 0.39 N/m, *β* = 0.42 for the MDCK II cell compressed with 0.5 μm/s and *T*_0_ = 0.57 mN/m, *K*_A_ = 0.24 N/m, *β* = 0.43 for the same cell compressed at 5 μm/s. F) Compression-relaxation curve of a MDCK II cell after blebbistatin treatment showing substantial softening. Blue line: Hertz fit (*E*_0_ = 62 Pa, *β* = 0.41). Red line: Evans fit (*T*_0_ = 0.02 mN/m, *K*_A_ = 0.002 N/m, *β* = 0.57) G) MDCK II cell subjected to calyculin A treatment that increases motor activity. Blue line: Hertz fit (*E*_0_ = 1097 Pa, *β* = 0.18). Red line: Evans fit (*T*_0_ = 1.6 mN/m, *K*_A_ = 2.36 N/m, *β* = 0.4).

We used an atomic force microscope to examine the viscoelastic properties of fibroblasts (3T3) and MDCK II cells in a confluent and suspended state. For parallel-plate compression experiments, tipless cantilevers were used to compress weakly adhering cells (Fig. 1), while cantilevers equipped with spherical (diameter: 3.5, 6.6, 15 μm) and conical tips (18° half cone angle) were employed for indentation experiments. We used constant approach and retraction velocities between 0.5 – 25 μm/s and a relaxation time of several seconds. As indicated, either blebbistatin or calyculin A were added to cell medium shortly before cell seeding. Detailed descriptions can be found in the SI.

Fig. 1D shows a typical compression-relaxation experiment of a single MDCK II cell using parallel-plate geometry. It is divided into the compression phase (i) during which the cell is loaded at constant velocity until the yield force is reached at *t*_m_ and subsequent force relaxation (ii) at constant distance between the plates. The full curve from the contact to the end of the relaxation curve was modeled with eq. (2) by adapting the three fitting parameters (red line), cortical tension *T*_0_, area compressibility modulus *K*_A_, and the power law exponent *β*. Eq. (2) requires input regarding the size of each cell (*R*_c_), which was measured using light microscopy prior to compression. For comparison, we also fitted the viscoelastic Hertz model (blue lines) to the data, which falls short in describing the curves, especially at low strain where tension dominates, and directly at the onset of relaxation. As a consequence, *β* values obtained from viscoelastic Hertz mechanics are systematically smaller than those provided by the Evans model (see SI). Fig. 1E shows representative fits of eq. (2) to compression-relaxation curves of MDCK II cells loaded with 0.5 μm/s and 5 μm/s, respectively. As required, the viscoelastic parameters are not impacted in this moderate velocity regime. However, since hydrodynamic drag at the onset of the compression curve is only negligible at low approach speed, subsequent experiments were carried out predominately at low speed (≤1 μm/s). The impact of blebbistatin and calyculin A on the compression-relaxation curves is shown exemplarily in Fig. 1F/G, while mean values are provided in Fig. 3. The softening of cells due to stalling of myosin motors is mirrored in smaller cortical tension and larger power law exponents, compared to untreated cells. This is particularly distinct for MDCK II cells, while fibroblasts in suspension are less affected (see SI). An increase in *β* is indicative of cortex fluidization, which we attribute to a loss of transient cross-links otherwise provided by myosin bundles. Administration of calyculin A, which is a phosphatase inhibitor that increases myosin II activity, generates only slightly larger pre-stress (contractility) and smaller *β* values, indicative of cell stiffening. Here, the drug turned out to be mildly toxic obscuring the effect of enhanced contractility. Knowledge of cortex thickness and mesh size (see SI, Fig. 1B/C), allows to estimate the area compressibility modulus from 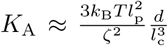 with the distance between cross-links 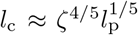 and the persistence length *l*_p_ of 17 μm [19]. With a mesh size of *ζ* = 25 - 150 nm and a cortex thickness *d* in the range of 100-1500 nm we arrive for *K*_A_ at values between 0.3 mN/m up to 11 N/m, which is in good accordance with our results. Notably, the same arguments leave us with Young’s modules in the range of 3 kPa up to 7 MPa, which is at least two orders of magnitude higher than values obtained from fits of the Hertz model (Fig. 2). Experimentally, the validity of a viscoelastic model can be verified by testing whether the two models generate viscoelastic parameters that are independent of the choice of indenter geometry or size. For this purpose, we used confluent MDCK II cells, which are easily probed with different indenter geometries and adapted the model according to the new overall geometry (Fig. 2, SI). Generally, confluent MDCK II cells are softer than those in suspension. Importantly, we find that the Young’s modulus obtained from the Hertz model depends on the size of the indenter. Larger radii of spherical and conical indenters result in systematically smaller Young’s modules. In contrast, neither cortex tension nor area compressibility modules depend on the indenter size rendering the Hertz model unsuitable to provide geometry-invariant viscoelastic parameters. The Evans model, however, is suitable to describe force relaxation curves over the entire experimental timescale independent of compression speed and indenter geometry. A poroelastic behavior of the cells (MDCK II) was proposed to describe the initial relaxation response after fast loading [22]. Here, we show that this initial drop is well captured by simple power law rheology but requires treatment of the actomyosin cortex as a pre-stressed shell. Therefore, the Evans model paves the way to address a fundamental problem in cell rheology, the apparent discrepancy between the rheology of living cells with glassy dynamics providing *β* values of 0.2 and the rheology of transiently cross-linked actin networks expecting *β* values of 0.5 reflecting the broad distribution of relaxation times [15]. Firstly, we found that the power law exponents obtained from the Hertz model are systematically lower 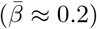 than those from the Evans model 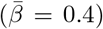. Secondly, cells with stiffer cortices display a smaller power law exponent (Fig. 3). Specifically, *β* decreases for both cell types logarithmically with *K*_A_ (Fig. 3B, SI). The same behavior has also been predicted by Gardel *et al.* [23] for the differential modulus and Kollmannsberger *et al.* [14] for the compliance of various cell types. Importantly, the model also allows to correlate internal pre-stress with fluidity. We found that an increase in internal stress is accompanied by a reduction of the power law coefficient (Fig. 3A) suggesting that cells with a stiffer, more contractile cortex are also less fluid. We also artificially increased tension by addition of glutardialdehyde (GDA) that contracts the cortex to generate solid-like shells with extremely low *β* values (Fig. 3). Recently, Mulla *et al.* found that artificial reversibly cross-linked actin networks show a decrease of *β* with increasing stress [16] suggesting that the glassy dynamics of the cortex are a natural consequence of transient cross-links combined with intrinsic pre-stress. Here, we can confirm that the source of the pre-stress are indeed myosin motors. In the absence of motor activity and therefore low pre-stress *T*_0_, *β* is close to 0.5, as expected for reversibly cross-linked actin filaments [15]. Notably, Yao et al. examined the rheology of actin networks cross-linked by *α*-actinin showing that external stress delays the onset of relaxation and flow, essentially extending the regime of solid-like behavior to much lower frequencies [24]. This was attributed to a catch-bond behavior of cross-linkers.

**FIG. 2.**
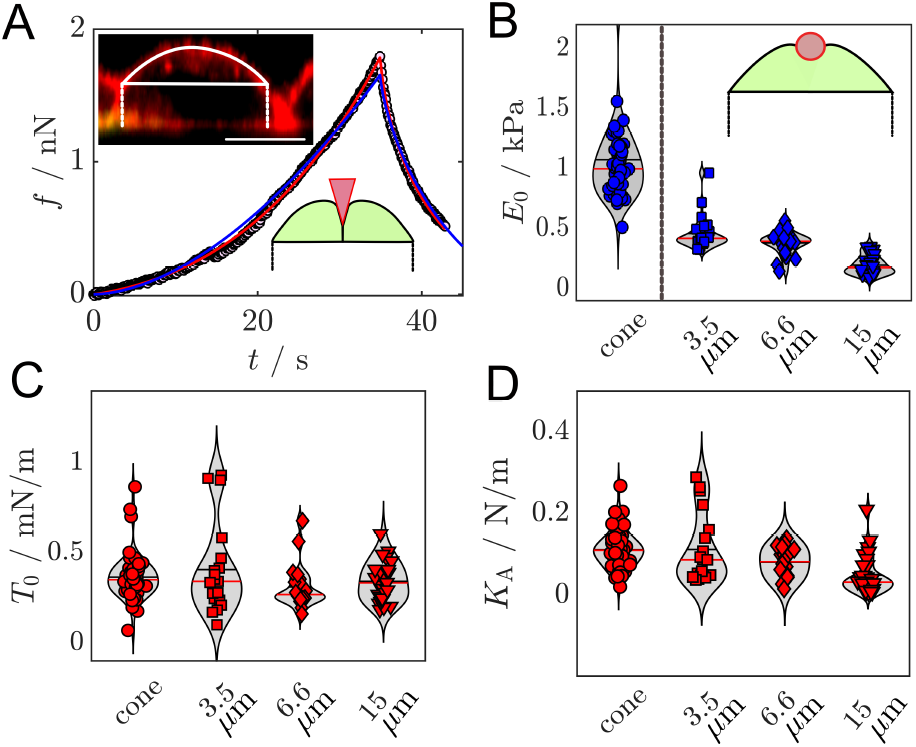
A) Indentation-retraction curve of a confluent MDCK II cell probed by a conical indenter and subject to fitting with the viscoelastic Evans model (red line) and Hertz model (blue line). The insets show a cross section of the cell and the computed shape after indentation, respectively (scale bar: 10 μm). B) Young’s modulus *E*_0_ of confluent MDCK II cells obtained from fitting a viscoelastic Hertz model to indentation-relaxation experiments performed with different indenter geometries (cone, spheres with various diameters). C/D) *T*_0_ and *K*_A_ of the same cells as in B) obtained from fitting with the viscoelastic Evans model.

**FIG. 3.**
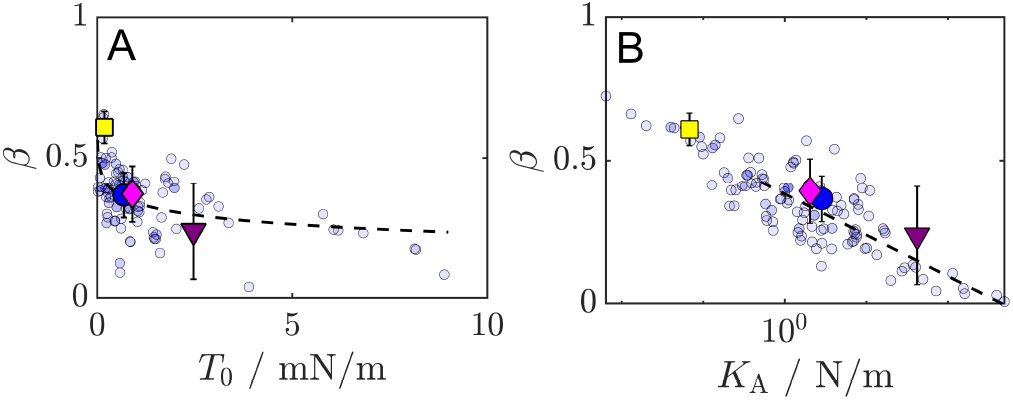
Power law exponents *β* of MDCK II cells as a function of pre-stress *T*_0_ and area compressibility modulus *K*_A_. Yellow square: mean value for blebbistatin-treated cells; blue circle: untreated cells; magenta circle: calyculin A-treated cells; purple triangle: GDA-fixated cells. Dotted lines represent fits illustrating the predicted logarithmic dependence of *β* on the elastic modules and the internal pre-stress [14].

In conclusion, we found that a viscoelastic shell model is capable of describing cell compression and relaxation experiments over the entire time scale in a consistent manner. MDCK II cells show a decrease in cortex fluidity with increasing pre-stress, thereby closing the gap between rheological experiments of artificial actin networks and living cells.

The work was financially supported by the DFG (SFB937(A8): AJ and MT; SFB 1027(A9): FL) and the VW foundation (‘Living Foams’: AJ and MT).

